# Age-specific gene signatures underlying the transcriptomes and functional connectomes of human cerebral cortex

**DOI:** 10.1101/2020.09.15.297754

**Authors:** Xingzhong Zhao, Jingqi Chen, Peipei Xiao, Jianfeng Feng, Ning Qing, Xing-Ming Zhao

**Affiliations:** Institute of Science and Technology for Brain-Inspired Intelligence, Fudan University, Shanghai 200433, China; Key Laboratory of Computational Neuroscience and Brain-Inspired Intelligence, Ministry of Education, China; Department of Computer Science, Tongji University, Shanghai 200092, China; Department of Computer Science, University of Warwick, Coventry, UK; Collaborative Innovation Center for Brain Science, Fudan University, Shanghai, China; Department of Mathematics, University of California, Irvine, CA, 92697, USA; School of Mathematical Sciences, Fudan University, Shanghai, China

## Abstract

The human cerebral cortex undergoes profound structural and functional dynamic variations across the lifespan, whereas the underlying molecular mechanisms remain unclear. Here, with a novel method TCA (Transcriptome-connectome Correlation Analysis), which integrates the brain functional MR magnetic resonance images and region-specific transcriptomes, we identify age-specific cortex (ASC) gene signatures for adolescence, early adulthood, and late adulthood. The ASC gene signatures are significantly correlated with the cortical thickness (*P*-value <2.00e-3) and myelination (*P*-value <1.00e-3), two key brain structural features that vary in accordance with brain development. In addition to the molecular underpinning of age-related brain functions, the ASC gene signatures allow delineation of the molecular mechanisms of neuropsychiatric disorders, such as the regulation between *ARNT2* and its target gene *ETF1* involved in Schizophrenia. We further validate the ASC gene signatures with published gene sets associated with the adult cortex, and confirm the robustness of TCA on other brain image datasets.

## Introduction

The cerebral cortex, making up over three quarters of the human brain volume, plays key roles in the cognition system(*1*). During maturation and aging, the cerebral cortex was suggested to exhibit dynamic functional connectivity (*2, 3*). For example, with aging, the long-range functional connectivity decreases among medial prefrontal cortex (mPFC), posterior cingulate (pC) and lateral parietal cortex (LP), while the connectivity among the left supplementary motor area, the right inferior temporal gyrus and the left temporal pole increase significantly, moreover, this regionally heterogeneous age effects were mainly detected in several functional sub-networks (e.g. default mode network (DMN)) (*3, 4*). The aberrance of either functional connectivity was found to lead to neuropsychiatric disorders(*5*). For example, children with Autism Spectrum Disorder (ASD) show reduced short and long-distance connectivity, especially in default and higher-order visual areas (*6*).

Much effort has been made to explore the molecular mechanisms underlying the structural or functional variations of the cerebral cortex. Using transcriptomic analysis, the glucocorticoid receptor gene *NR3C1* has been found correlated with cortical thinning during adolescence, indicating its important role in cortical maturation (*7*). In addition to being correlated with structural variations, using the correlation between the dynamic patterns of gene expression and spatial topography of group difference in some imaging phenotype, could provide molecular mechanisms that underlie functional connectivity of brain cortex (*8*). By analyzing gene expressions using Allen Human Brain Atlas, Hawrylycz *et al*. identified a set of 136 genes whose co-expressions were strongly correlated with the functional connectome built over healthy adolescents, implying their roles of neuron projection and axon guidance (*9*). Anderson *et al*. found strong associations between another set of genes and the limbic functional networks based on transcriptome data, and showed that some of them were enriched in the inhibitory interneuron marker somatostatin receptor pathway and they were potential risk genes for psychiatric disorders (*10*). Nevertheless, existing studies often focus on a specific circuitry or an age group, the molecular dynamics or changes across the lifespan for cerebral cortex remain largely unknown.

Here we present a method, TCA (Transcriptome-Connectome correlation Analysis), to identify gene signatures associated with cortex development and functionality across lifespan. By integrating age-specific brain connectomes with gene expression data, we identify gene signatures for three distinct age groups, i.e. adolescence (aged 8-20), early adulthood (aged 20-40) and late adulthood (aged 40-82). We show that these gene signatures can explain age-specific functional dynamics of cerebral cortex during maturation and aging, and may help understand the molecular mechanisms of neuropsychiatric disorders.

## Results

### Age-specific gene signatures illustrate trajectories of cortical maturation and aging

As shown in Fig.1, we presented an approach, TCA (Connectome-transcriptome Correlation Analysis), to identify the age-specific cortex (ASC) genes based on the integration of fMRI connectomes and gene connectivity maps, which derived from the fMRI and transcriptome data across multiple cortical regions for each subject. The mean values of the connectomes and gene connectivity maps for each age group were used to perform the correlation analysis (Materials and Methods). As a result, three sets of age-specific cortex (ASC) genes were identified consisting of 155, 114, and 136 genes for adolescence (aged 8-20), early adulthood (aged 20-40), and late adulthood (aged 40-82), respectively (Tables S1 and S2). Most of them (90.5%) only showed up in one signature, indicating the age-specificity of these signatures (Fig. S1). One signature gene, *MYLK* (encoding myosin light chain kinase), was observed in all the three age groups. *MYLK* has been reported to induce retraction of mature oligodendrocyte processes, and this process plays an important role in myelin formation and maturation in the central nervous system (*11*).

**Fig.1.**
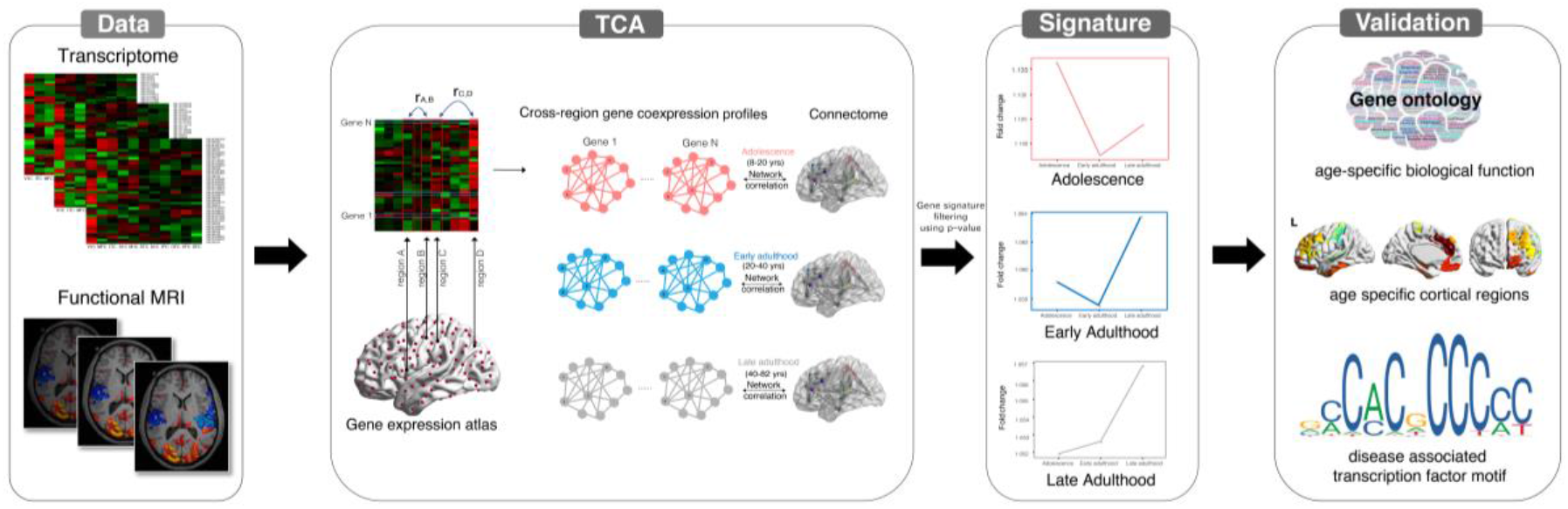
Overview of the connectome-transcriptome correlation analysis. The first column shows the input data for TCA, include the functional MR images and transcriptome data. The second column depicts the main idea of TCA, calculating the correlation between the functional magnetic resonance imaging(fMRI) connectome and the connectivity map (coexpression profile)) for each gene (Methods). The third column shows the expression patterns of the age-specific cortex (ASC) gene signatures identified by TCA for each age group. The last column illustrates the functional validation for the roles of ASC gene in cortex maturation and aging, through identifying the age-specific biological processes, the correlation between gene expression and brain physical structural measures (e.g. myelination and thickness), and the relevant molecular details associated with brain disorders.

By examining their expression dynamics, we noticed that the signatures demonstrated brain region-specific and age-specific expressional patterns. Compared with other genes in the genome, the genes from the three signatures were significantly highly expressed (T-test *P*-values = 1.96e-10, 2.55e-10, and 4.97e-9 for adolescence, early adulthood, and late adulthood, respectively), where the ASC genes for early adulthood were most highly expressed, followed by those for adolescence and for late adulthood (Fig. 2A and Fig. S2). We then identified the brain regions in which each signature were highly expressed, and referred them as “highlighted regions” for the corresponding age group (Fig. 2B, see Materials and Methods). Three brain regions that were found to be highlighted for adolescence, including dorsolateral prefrontal cortex (DFC), primary somatosensory cortex (S1C), and ventrolateral prefrontal cortex (VFC). Previous, both DFC and VFC have been reported to be involved in the process of ‘executive’ functions (e.g. working memory and response inhibition), which were highly demanded during adolescence (*12, 13*). For the early adulthood group, the highlighted brain regions include medial prefrontal cortex (MFC), orbital prefrontal cortex (OFC), and inferior temporal cortex (ITC), which were reported to be correlated with cognitive, behavioral, and emotional control (*14*). For the late adulthood group, the functional decline in some basic cognitive areas of the frontal lobe has been reported (*15*). Consistent with this prior knowledge, the regions VFC, MFC, ITC, and OFC were highlighted by the ASC genes for late adulthood, and these brain regions were known to be responsible for memory, information processing, and the ability for language (*16–18*).

**Fig. 2.**
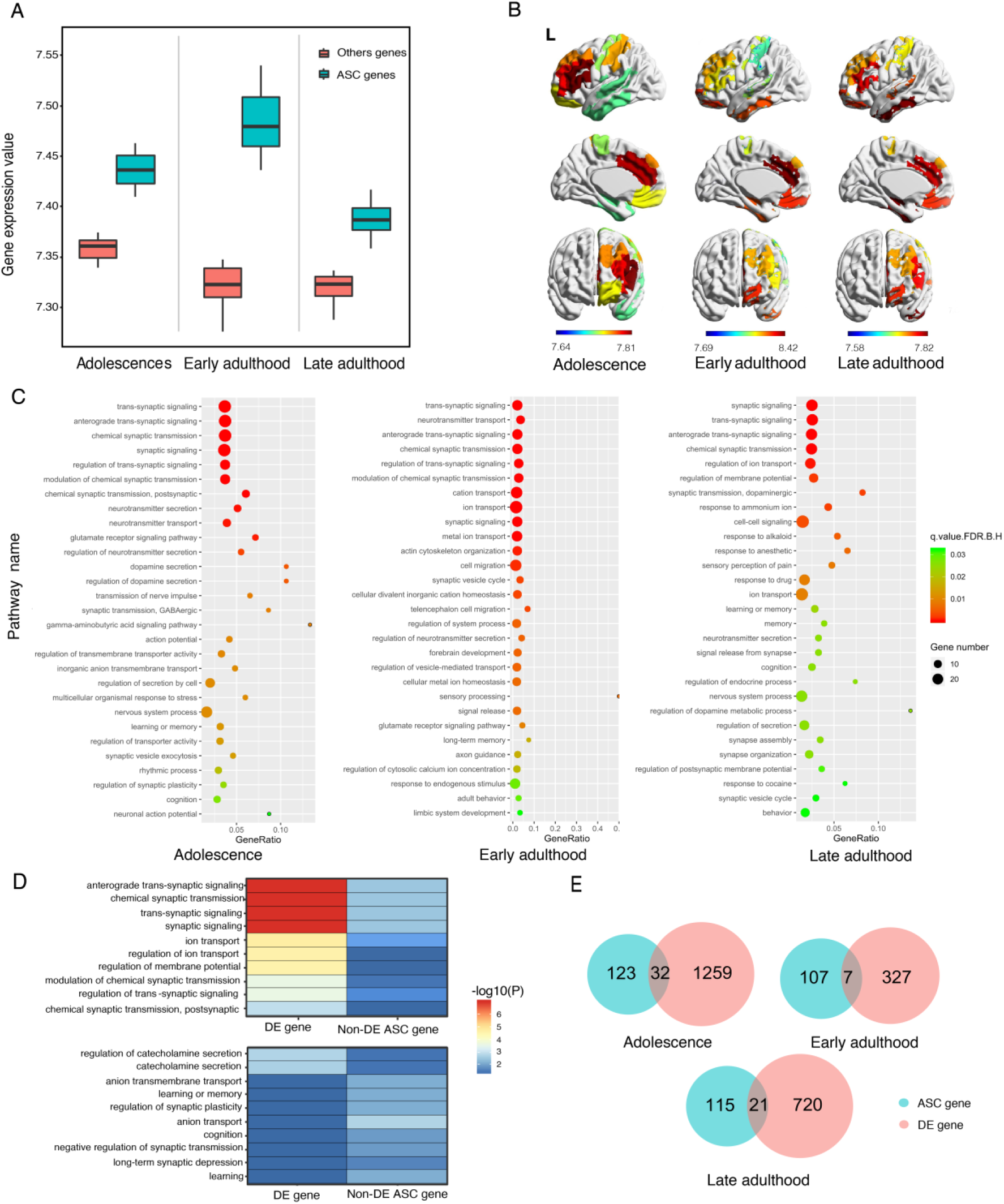
The brain regions and biological processes highlighted by the ASC genes. **(A)**, Comparison of the expression level of the ASC genes identified by TCA against that of other genes (all background gene except ASC genes) at different age stage. (**B)**, Cortical regions highlighted by the ASC genes for each age group. The highlighted regions mean that the ASC gene were highly expressed. (**C)**, The biological processes enriched by the ASC genes for each age group. (**D**), The enriched biological processes that are different for the ASC genes and the differentially expressed genes at adolescence. Colors represent enrichment *P*-values of the bilogical processes, where the darkest blue bars refer to no enrichment. (E), The overlap between the ASC genes and the differentially expressed genes for three age groups.

Together, the identified age specific gene signatures showed specific spatiotemporal patterns, highlighting brain regions that are important for brain functions during the corresponding age periods.

### Age-specific gene signatures characterize age-relevant cortical functions

Next, we investigated the biological functions of the three ASC gene signatures based on the annotations from Gene Ontology (GO, http://geneontology.org). We studied a selection of significantly enriched biological processes for each age group (Fig. 2C, *FDR* ⩽ 0.05, Materials and Methods). As expected, some basic functions important for neuron development and cortex structure were enriched across all three age groups, e.g. chemical synaptic transmission, signal release, and calcium ion binding. For each age group, the enriched functions of the corresponding ASC genes were generally consistent with prior knowledge of cortex development and functionality around that age period (Fig. 2C and Table S3). For example, the ASC adolescent genes were enriched with the processes related to neuron and synapse development, e.g. neurotransmitter receptor activity (GO:0030594) and learning or memory (GO:0007611), which were obviously important for the adolescent brain. Specifically, the ASC adolescent genes *PPFIBP1* and *ARL15* have been reported to participate in the guidance and development of axons, and *GRIA1*, a subunit of the AMPA receptor, have been found to associated with mature synaptic function (*19*),(*20*). The ASC genes for early adulthood were enriched with processes involving motion control and cell differentiation, e.g. limbic system development (GO:0005157) and cerebral cortex cell migration (GO:0021795), and those for late adulthood were enriched with immune cells and inflammation-related biological processes such as macrophage colony-stimulating factor receptor (GO:0005157).

Conventionally, age-specific genes have been identified with differential expression analysis between distinct age groups (*21–23*). Compared with genes differentially expressed between the three age groups considered here, some of the ASC genes were also differentially expressed genes (DEGs) (20.6%, 6.1%, and 15.4% for adolescence, early adulthood, and late adulthood, respectively; Fig. 3E, Table S4). The functional enrichment analysis shows that both DEGs and ASC genes were enriched in common neuronal functions, e.g. synaptic signaling (GO:0099536) (Table S5). However, the ASC genes not differential expressed were found enriched with age-specific advanced cognitive functions, which were missed by DEGs (Fig. 3D). For example, the non-DE ASC genes for adolescence were enriched with learning, memory, and cognition, while the non-DE ASC genes for early adulthood were enriched with forebrain development. Therefore, our identified age-specific gene signatures were able to characterize the age-specific cortical functions better than DEGs.

**Fig. 3.**
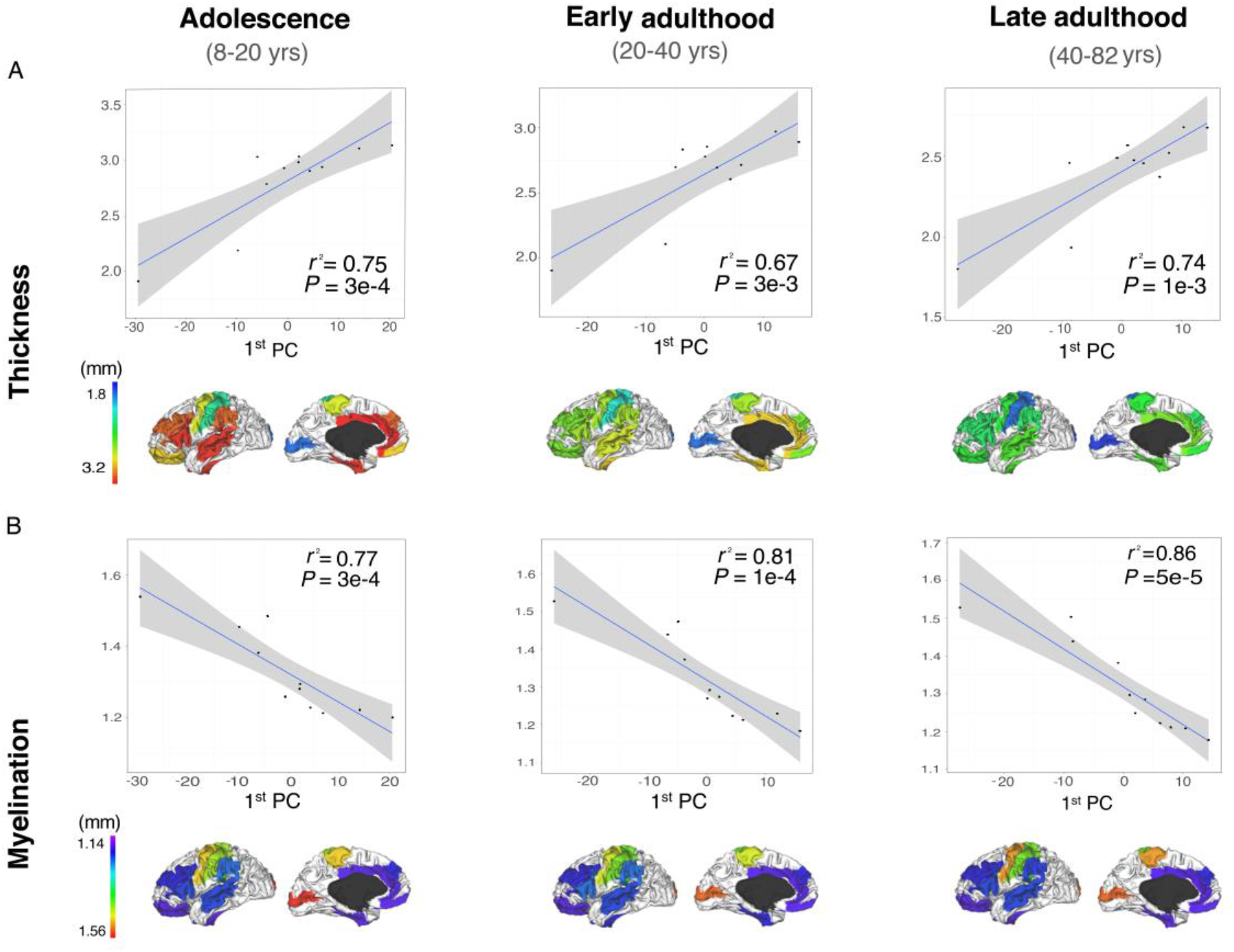
Age-specific cortical structural features significantly associated with the ASC genes. Correlations between the expression levels of the ASC genes and cortical thickness (top) and myelination (bottom) were calculated as the Pearson correlation coefficient (*r^2^*) between the first principal component (1^st^ PC) of the expression of the ASC genes and the cortical structural measures (for thickness or myelination) for each age group. *P* denotes the *P*-value for the Pearson correlation test.

**Fig. 4.**
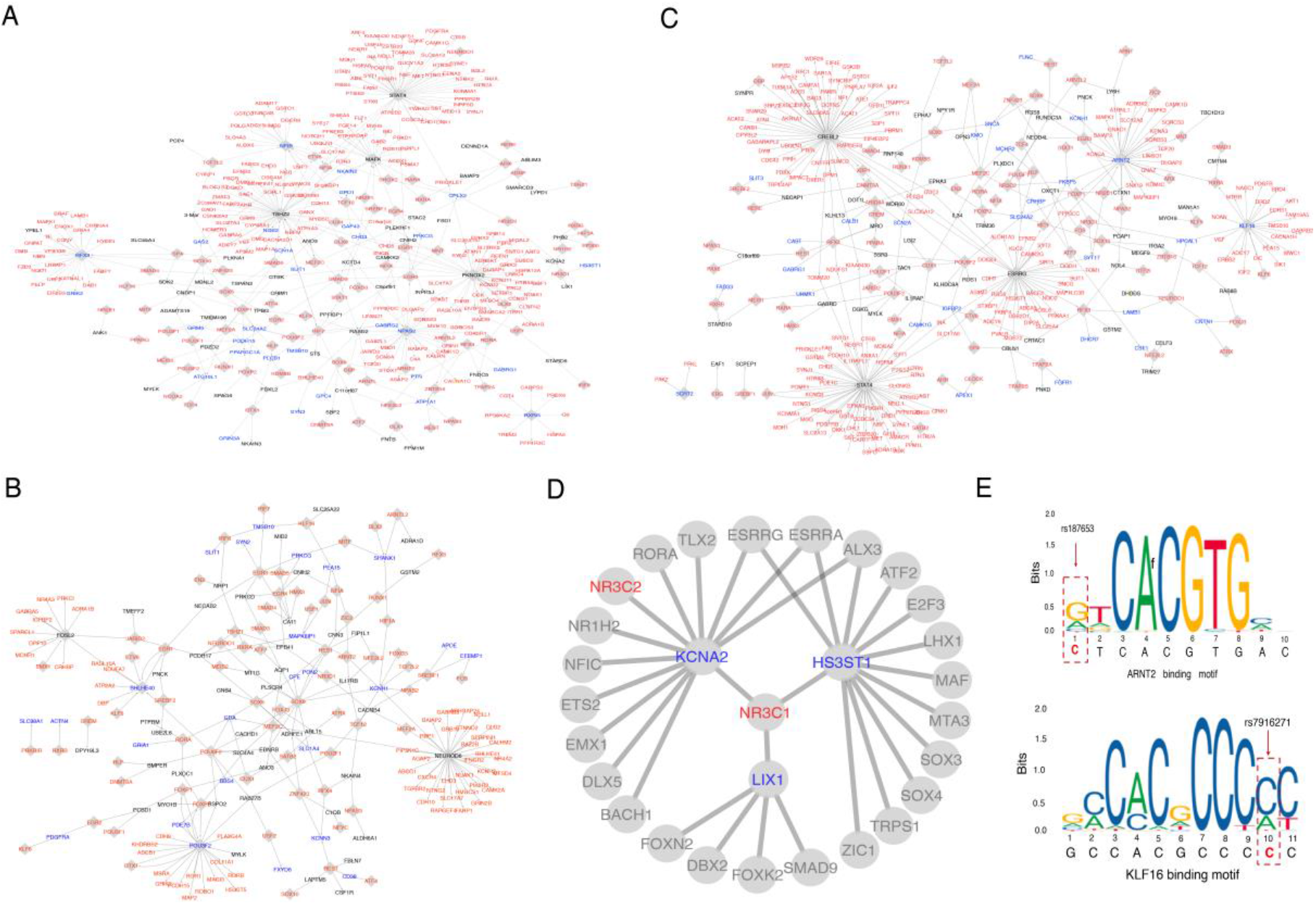
The role of ASC genes in neuropsychiatric disorders. **(A,B, C)**, The disease sub-TRNs for adolescence, early adulthood and late adulthood, featuring only the regulatory relationships between the ASC genes and neuropsychiatric disease genes. Nodes with black labels are the ASC genes, and nodes with red labels are disease gene. If a node is both an ASC gene and a disease gene, it is labeled in blue. The size of each node represents its degree in the sub network. (**D),** The ASC genes regulated by their upstream regulator — the TF *NR3C1* (in the middle, red labeled, indicating it was a disease gene for ADHD), as well as the other TFs regulated ASC gene, in the sub-TRN for adolescence. The nodes with blue label are the ASC genes, other TFs of the ASC genes that are disease genes for ADHD are labeled in red, while others are labeled in black. **(E)**, Disruption of the *KLF16* binding motifs by SNP rs7916271, and disruption of the ARNT2 binding motifs by SNP rs187653.

### Age-specific gene signatures are implicated in cortical structural dynamics

In previous studies, it has been indicated that morphological features of the cerebral cortex varied with age (*24*). For example, the cortical volume and thickness would change as cortex matures from childhood to adolescence, and these dynamic structural variations were closely associated with cognitive development (*25, 26*). Here, with the age-specific signatures, we wanted to see whether these ASC genes can provide an extra layer of explanation for age-related brain structural dynamics. For this purpose, we extracted structural features from structure MRIs of the same samples used in this study, including cortical volume, cortical surface area, cortical thickness, curvature, and myelination. With the first principal component of the expression profile of each ASC gene signature, we investigated the correlation between each pair of structural features and ASC signature across the 11 cortical regions. As shown in Fig. 3, the expression of ASC genes was significantly correlated with cortical thickness (*P*-values ranging from 3.00e-4 to 2.00e-3) and myelination (*P*-values ranging from 2.00e-4 to 1.00e-3), both of which were important for cortical development and cognitive abilities (*27, 28*). Take *GRIA1* as an example, it was one of the ASC genes for adolescence, and its expression was significantly correlated with cortical thickness (*P*-value = 8.14e-4). *GRIA1* encoded a subunit of the AMPA-selective glutamate receptor 1, while the metabolism of glutamate has been reported to play important roles in cortical thickness (*29*). Furthermore, we wanted to check whether our ASC genes overlap with gene sets previous reported to be associated with brain structural features. For example, a set of 62 genes have been reported significantly associated with cortical volume based on their expression, and we found that our ASC genes were significantly enriched in (*P*-value = 9.46e-8, Fisher exact test) and highly functionally similar to (GOSemSim (*30*) = 0.86) those 62 genes (*31*). These findings indicate that our gene signatures may provide molecular insights into structural variations during cortical maturation and aging.

### Age-specific gene signatures play central roles in age-specific brain transcriptional-regulatory networks

Recently, Pearl et al. have constructed a brain-specific transcriptional-regulatory network (TRN) by integrating sequence features and DNase footprinting data of the brain, further filtered by gene expression information in multiple brain regions (*32*). The full network consisted of 741 transcription factors (TFs) and 11,092 target genes, where 17 of our ASC genes were found to be among the 741 TFs. Taking advantage of this brain TRN, for each age group, we first extracted a sub-TRN where the expression levels of each pair of nodes (a TF and a target gene) connected by an edge were significantly correlated in that age group (Pearson correlation test, *FDR* ⩽ 0.05, Table S12). As a result, the sub-TRN for adolescence consisted of 479 TFs and 9,137 target genes, the sub-TRN for early adulthood consisted of 477 TFs and 8,953 target genes, and the sub-TRN for late adulthood consisted of 480 TFs and 8,941 target gene. In the three age-specific sub-TRNs, we examined the network-based properties of our ASC genes. Interestingly, in all three sub-TRNs, the degrees of ASC genes were significantly higher than others genes (*P*-values < 2e-16, Fig. S3), indicating that the ASC genes might have more central transcriptional regulatory roles in the brain. For example, NFIB, which was an ASC TF for adolescence and regulated 60 target genes in the sub-TRN of adolescence, has been reported essential for late fetal forebrain development and loss of NFIB gene leads to defects in basilar pons formation and hippocampus development (*33*). On the other hand, the target genes regulated by ASC genes in those sub-TRNs were significantly enriched in genes associated with neuropsychiatric disorders (Fig. S4). For adolescence, the target genes were significantly enriched in autism spectrum disorders (ASD, *P*-value = 1.66e-4) and schizophrenia (SCZ, *P*-value = 3.12e-4). The target genes of early adulthood were significantly enriched in depression (*P*-value = 4.79e-3) and bipolar(*P*-value=3.16e-5). For late adulthood, the target genes were not significantly enriched in any disorders.

Together, the ASC genes were found to play central regulatory roles in the brain transcriptional regulatory network, which may explain their important functional roles in cortex maturation and aging.

### Age-specific gene signatures are validated with independent datasets

In order to validate our ASC gene signatures, we compared our ASC gene signatures with two previously published gene sets for the adult brain—the RichSet(*34*) and MichSet(*9*) (see Materials and Methods). The RichSet contains 136 genes significantly correlated with cortex connectome of adults aged between 20 and 60, and the MichSet contains 75 genes defined in the similar way as RichSet for adults aged between 20 and 60. From Table 1, both RichSet and MichSet were significantly enriched in the ASC genes for age groups after adolescence, with *P*-values of 5.73e-12 and 7.83e-9, respectively (Fisher exact test). In addition to analyzing the overlap between gene sets, we examined the functional similarity between gene sets with the assumption that functional similar gene sets may have equivalent functional roles. Our ASC signatures were found to be highly functionally similar to both RichSet and MichSet, and the functional similarities between two gene sets were quantified by GOSemSim(*30*) (see Materials and Methods).

**Table 1.** Validation with known gene sets and test image datasets

|  | Gene sets |  | Image datasets |  |
| --- | --- | --- | --- | --- |
|  | RichSet(34) | MichSet(9) | IMAGEN | HCPS900 |
| **Age distribution** | 20-60 | 20-60 | 17,19 | 20-40 |
| **Gene number** | 136 | 75 | 46 | 129 |
| **Functional similarity with the ASC genes** | 0.84 | 0.88 | 0.76 | 0.75 |
| **P-value for the overlap with the ASC genes** | 5.73e-12 | 7.83e-9 | 1.20e-6 | 2.2e-16 |

We further tested the robustness of our TCA method, by replicating the computational process on the same transcriptome but two test image datasets—the resting-state fMRI data from the IMAGEN project(*35*) and the HCP S900 dataset(*36*) (see Materials and Methods). The two datasets were respectively collected from adolescent and early adult subjects (see Table 1). As a result, 46 genes (the IMAGEN dataset) and 129 genes (the HCP S900 dataset) were obtained. We noticed that these two gene sets were significantly enriched with the union of our ASC signatures for adolescence and early adulthood (*P*-value = 1.20e-6 and *P*-value < 2.2e-16). By examining further the functional similarities between the two gene sets with our ASC signatures, we found that both genes sets were functionally similar to the ASC signature with the similarity value larger than 0.70 (by GOSemSim (*30*)). The analysis demonstrates the robustness of our method, and indicate that TCA can be applied to datasets for other biological systems or contexts.

## Discussion

Both the transcriptome and the connectome of the human cerebral cortex undergo extensive changes across the whole lifespan. Here, with a novel method named Connectome-transcriptome Correlation Analysis (TCA), we identify age-specific cortex (ASC) gene signatures for three different age groups, i.e. adolescence, early adulthood, and late adulthood. These ASC genes are functionally enriched with cortex-specific biological processes, and their expression is significantly correlated with chronological structural changes of the cortex. These genes also play central roles in the transcriptional-regulatory network of the brain, and the disrupted regulations of them may contribute to the etiology of neuropsychiatric disorders. Taken together, through integratively analyzing transcriptome and connectome, we have successfully identified three age-specific gene signatures, which recapitulate the structural and functional dynamics during cortical maturation and aging, and pinpoint the molecular mechanisms underlying neuropsychiatric disorders.

Although the ASC gene signatures are identified with functional connectome instead of structural MRIs, the expression of our identified ASC genes is significantly correlated with cortical thickness and myelination, the two brain structural features known to vary with age. In fact, some genes reported to be correlated with brain structure are also contained in our ASC genes. For example, *SLIT3* has been reported to be associated with cortical volume (*37*) and *WNT3* with myelination (*38*), while both genes belong to the ASC genes, implying that the ASC genes are indeed correlated with brain structures. Therefore, the ASC gene signatures may help better capture the molecular mechanisms involved with the interplay between brain structure and function.

Recently, a growing body of evidence has suggested that many neuropsychiatric disorders are associated with certain genetic mutations. However, it is largely unclear how these genetic mutations contribute to the disorders. Moreover, these mutations themselves only account for part of the heritability of the neuropsychiatric disorders (*39*). Understanding the biology underlying the normal trajectories of cortical maturation and aging, as a result, may provide a global picture for how the disorder-associated mutations fit into the pathological process. Thus, our ASC genes may help interpret the mechanisms of neuropsychiatric disorders in at least the following three ways: 1) given that the ASC genes depict age-specific biological functions of the cortex, the regulatory relationships between the ASC genes and other genes that are disrupted by mutations may explain the functional impact of the mutations, as we have shown in this manuscript; 2) given the age-specific association between the ASC genes and the cortical functional and structural MRI features, disrupting the regulations of the ASC genes by the disorder-associated mutations can be mapped to a network of regional neuro-circuitries, such as the default-mode network (DMN) implicated in multiple neuropsychiatric disorders (*40*); 3) the ASC genes can also be presented as a set of candidate biomarkers whose disruptions may lead to disorder status.

In this work, we only focus on 11 regions of the cerebral cortex due to limited availability of the brain regional transcriptome data. These 11 regions may not precisely depict the functional details of cortex, as they are relatively broad regions. In addition, the relatively small sample size used may reduce the statistical power of our approach. With more and more brain transcriptome and imaging data released in the future, we may be able to obtain results with better reliability and higher resolution. Nevertheless, we have partially validated our results with previously identified gene sets, proving the robustness of our approach to some extent. It is also possible to incorporate other types of brain datasets into the TCA method, so as to gain insights from additional levels. As such, we present our method and the ASC genes to help understand the mechanisms underlying cortical maturation and aging. We hope our findings will be of value for the study of both the normal and the disease states of the brain.

## Materials and Methods

### Preprocessing of MRI data

The functional and structural MRI data used in this study are downloaded from the Human Connectome Project (HCP) Lifespan Pilot Project (http://lifespan.humanconnectome.org/). The dataset contains both functional and structural MRI images covering the whole brain, acquired from 27 subjects (aged 8-75). The subjects are divided into three age groups (8-20, 20-40, and 40-75; see details in Table S8). Both the structural and functional MRI images are preprocessed by following the HCP Pipelines with the default parameters (http://www.humanconnectome.org/documentation/HCP-pipelines). Then we adopt the surface-based cortical parcellations atlas of the Brodmann Area Map (Brodmann lh,colin.R via pals_R-to-fs_LR) to divide the left-hemisphere cortex into 41 spatially contiguous parcels as in the VDG11b parcellation of the Brodmann Area Map (*41, 42*). The HCP software, called Workbench Command (https://www.humanconnectome.org/software/workbench-command), is used to extract for each parcel of the BOLD time series signal across 420 time points and the structural metrics, including cortical volume, cortical surface area, cortical thickness, curvature, myelination and sulcus.

### Preprocessing of gene expression data

The gene expression data for the human cerebral cortex used in this study is downloaded from the NCBI Gene Expression Omnibus under the accession number GSE25219 (Affymetrix Human Exon 1.0 ST Array)(*43*). This microarray data contains 17,565 probes across 11 areas of the neocortex (NCX, Table S9), with a total of 1,340 cortical and subcortical tissue samples ranging from prenatal stages to late adulthood. We preprocess the microarray data as described by Kang et al (*43*). Firstly, a stringent criterion (log2-transformed signal intensity >= 6 in at least one sample, and mean detection-above-background P-value <= 0.01 in at least one NCX region during at least one period) is used to define “expressed” probes, which results in 12,837 (73.1%) probes for further analysis. Secondly, we map the 12,837 expressed probes to 17,650 protein-coding genes based on the annotation of Affymetrix Human Exon 1.0 ST Array (as of February 18th, 2019). When multiple probes are mapped to the same gene, the mean value of the probes is used as the expression value of the gene. Finally, we extract the preprocessed gene expression profiles for subjects aged 8 to 78 (232 samples) and re-group them into three age groups (8-20, 20-40, and 40-78), in order to match the MRI datasets.

### Known gene signatures

Two gene sets, named as the RichSet (*34*) and MichSet (*9*), that have been reported as gene signatures of adult cortex are used for validation in this study. Both gene signatures are defined based on the gene expression data from the Allen Human Brain Atlas of six adults (20 to 60 years old). The RichSet containing 136 genes was generated by integrative analysis of gene expression data with connectome derived from 15 healthy right-handed subjects(*34*). The MichSet (75 genes) is generated based on gene expression with HCP S500 connectome data (*9*), where a subset of the genes whose expression were highly associated with functional connectivity are selected.

### Test image datasets

We download the connectome data from IMAGEN (https://imagen-europe.com) and HCP S900 project (www.humanconnectome.org/study/hcp-young-adult). The participants of the IMAGEN project are healthy adolescents aged 14 or 19, as described in Schumann *et al*. (*35*). We use the SPM12 (https://www.fil.ion.ucl.ac.uk/spm/software/spm12/) preprocessing the IMAGE rsfMRI images with realignment, slice-timing correction, movement correction, non-linear warping into MNI space using a custom EPI template, and Gaussian-smoothing at 5 mm full-width half-maximum. The HCP S900 project has 900 healthy participants aged from 22 to 35. We preprocess the HCP S900 dataset by following the HCP pipelines with default parameters (http://www.humanconnectome.org/documentation/HCP-pipelines). The parcellation of the brain is carried out as described above. We then extract the 11 cortical regions of left-hemisphere used in this study with the mean value of voxels in a ROI as the value for the ROI.

### Connectome-transcriptome Correlation Analysis

Firstly, we design a score *GS_score_* to describe the stability of gene expression among individuals. For each gene in each age group, the *GS_score_* is defined as the average of the Pearson’s Correlation Coefficients (PCC) between all pairs of subjects in this age group based on their gene expression profiles over 11 cortical regions as follows.

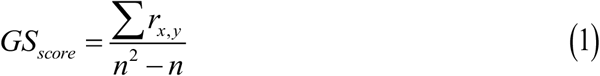

where ∑ *r_xy_* is the summary of pairwise PCCs between subjects (*x, y, x* ≠ *y*) for a specific gene in each age group, and *n* is the total number of subjects within this age group. For each age group, we rank all the genes based on their *GS_score_*, and select the top 10 percent genes as the stable genes for further analysis (Table S10). Secondly, for each stable gene in each age group, we construct a gene network where each node is a cortical region, and the weight accompanying each edge is the normalized co-expression correlation coefficient between a pair of regions across all the subjects in the age group as defined below.

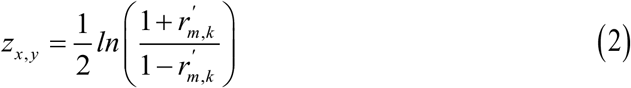

where 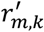 is the co-expression correlation coefficient between cortical region *m* and *k*. Thirdly, for each subject, a functional connectome is constructed from the BOLD time series data across 420 time points, where each node is a cortical region and the weight of each edge is the normalized correlation coefficient between a pair of cortex regions as defined in Eq (2). Then, the average of the connectomes of all subjects in each age group is used as the resultant functional connectome for the age group. Finally, for each age group, a list of genes whose connectivity are significantly correlated (*P*-value<=0.05) with the functional connectome of the same age group are kept for further analysis. To avoid false positives, the permutation test is adopted by randomly permutating sample labels, and the above mentioned computation process is run for 1000 times. For each gene, the *P*-value is defined as the ratio of the number of times among 1000 times that the gene has a higher correlation coefficient with the functional connectome compared with the original one. Consequently, for each age group, the list of genes with *P*-value less than 0.05 is defined as the age-specific cortical (ASC) gene signature for the corresponding age group.

### Risk genes of neuropsychiatric disorders

The Attention Deficit Hyperactivity Disorder (ADHD) related genes are acquired from the ADHDgene database (http://adhd.psych.ac.cn/), and we select the genes supported by at least 60% of all studies included in the database(*44*). Autism Spectrum Disorder (ASD) related genes are downloaded from the AutDB database (http://autism.mindspec.org/autdb/) and a latest published work (*45, 46*). Alzheimer’s disease related genes used in this paper are downloaded from the ALzGene database (http://www.alzgene.org/) (*47*). Schizophrenia risk genes are downloaded from the SZGene database (http://www.szgene.org/) and wang et. Al (*48, 49*). Bipolar risk genes are download from DisGeNet (*50*). Parkinson’s disease associated genes are downloaded from the PDGene database website (http://www.pdgene.org/) (*51*). Depression associated genes are downloaded from the website (http://www.polygenicpathways.co.uk/depression.htm). All these risk genes can be found in Table S11.

### Gene function annotation

The functional enrichment analysis on ASC genes is performed using the ToppGene portal (https://toppgene.cchmc.org/, latest update on 2019-8-21)(*52*). The ToppGene portal incorporates a comprehensive list of gene function annotation databases, and the P-value for a query gene list is obtained with a hypergeometric test, with various choices for multi-test correction. From the databases, we select Gene Ontology (GO) annotations (biological process, cellular component, and molecular function), pathway annotations (KEEG and REACTOME), and disease gene annotations (OMIM and GWAS), for functional enrichment analysis of the ASC gene sets.

### Differentially expressed genes

One-way Analysis of variance (ANOVA) is adopted to identify the differentially expressed (DE) genes among the three age groups, and do the post-hoc analysis by Tukey’s test to select gene sets that are specific in one group (Table S4).

### R functions used in the analysis

The analysis is performed in R with the following default functions: Pearson correlation coefficient - cor(); ANOVA - aov(); Tukey’s test -TukeyHSD(); Fisher’s exact test - fisher.test(); Principal component analysis (PCA) - prcomp(); nonlinear fitting for BOLD time series signals across all time points - loss(). Functional similarity between two sets of genes is carried out with functions from the R package “GOSemSim”(*30*).

### Identifying the potential neuropsychiatric disorders mechanism of TFs

Neuropsychiatric disorders associated single nucleotide polymorphisms (SNPs, *P*-value ≤ 5e-6) are collected from the summary statistics of Genome-Wide Association Studies (GWAS) hosted in the PGC data portal (http://www.med.unc.edu/pgc/) and the GWAS Catalog database (https://www.ebi.ac.uk/gwas/). We use the online tool LiftOver (http://genome.ucsc.edu/cgi-bin/hgLiftOver) to convert SNP locations from genome versions hg18 and hg19 to genome version hg38. Next, we download the genome annotation file from NCBI (hg38; ftp://ftp.ncbi.nih.gov/genomes/H_sapiens/GFF/), and extract the gene position information. We download the motif files of all human TFs from Jaspar (http://jaspar.genereg.net/)(*53*), and use FIMO (http://meme-suite.org/tools/fimo) to get TF binding motif information across the whole genome. Furthermore, we use BEDTools(*54*) to match the locations of risk SNPs, gene promoter regions (TSS ± 10KB), and the transcription factor binding motifs.

## Acknowledgments

We thank Dr Tianye Jia for his suggestions on the differential gene analysis.

## Author contributions

Xing-Ming Zhao conceived the project, designed the framework. Xingzhong Zhao and Peipei Xiao collected the data, and analyzed the data. Xingzhong Zhao wrote the manuscript. Jingqi Chen, Xing-Ming Zhao, Ning Qing, and Jianfeng Feng supervised the data analyses and revised the manuscript.

## Data and materials availability

All scripts are written in R. Scripts for the TCA method and related statistics result can be freely accessed at https://github.com/Soulnature/TCA. Additional data related to this paper may be requested from the authors.

